# Genetic encoding of targeted MRI contrast agents for *in vivo* tumor imaging

**DOI:** 10.1101/799411

**Authors:** Simone Schuerle, Maiko Furubayashi, Ava P. Soleimany, Tinotenda Gwisai, Wei Huang, Christopher Voigt, Sangeeta N. Bhatia

## Abstract

Tumor-selective contrast agents have the potential to aid in the diagnosis and treatment of cancer using noninvasive imaging modalities such as magnetic resonance imaging (MRI). Such contrast agents can consist of magnetic nanoparticles incorporating functionalities that respond to cues specific to tumor environments. Genetically engineering magnetotactic bacteria to display peptides has been investigated as a means to produce contrast agents that combine the robust image contrast effects of magnetosomes with transgenic targeting peptides displayed on their surface. This work reports the first use of magnetic nanoparticles that display genetically-encoded pH low insertion peptide (pHLIP), a long peptide intended to enhance MRI contrast by targeting the extracellular acidity associated with the tumors. To demonstrate the modularity of this versatile platform to incorporate diverse targeting ligands by genetic engineering, we also incorporated the cyclic αv integrin-binding peptide iRGD into separate magnetosomes. Specifically, we investigate their potential for enhanced binding and tumor imaging both *in vitro* and *in vivo*. Our experiments indicate that these tailored magnetosomes retain their magnetic properties, making them well-suited as T2 contrast agents, while exhibiting increased binding compared to wild-type magnetosomes.

## Introduction

Robust imaging of tumors for diagnosis and treatment monitoring has the potential to improve healthcare outcomes and save lives. Wide recognition of this principle, combined with established screening practices based on noninvasive imaging modalities, has spurred investigation into contrast agents that aid in distinguishing tumors from surrounding tissue.^1,2^ Among possible imaging modalities, magnetic resonance imaging (MRI) is especially appealing because magnetic fields are relatively innocuous compared to ionizing radiation, and targets throughout the body can be readily resolved.^3^ Contrast agents designed to identify tumors do so by exploiting their unique physical and biochemical characteristics. An effective tumor-selective imaging agent must therefore combine properties that make it robustly detectable via the imaging modality with features that lead to preferential accumulation or enhanced contrast effects in tumors. Because targeting strategies often rely upon biomarkers specific to particular types of tumors, tumor-selective contrast agents based on generalized characteristics of tumors are especially desirable.

MRI contrast agents typically function by detectably altering the longitudinal (T1) or transverse (T2) relaxation times of nearby hydrogen nuclei.^3^ Synthetic magnetic nanomaterials, especially ferrite nanoparticles, have been deployed as MRI contrast agents due to their magnetic properties and biocompatibility. For instance, size-dependent surface effects in ultrasmall ferrite nanoparticles have spurred their investigation as T1 contrast agents, offering a potential alternative to the paramagnetic gadolinium ion complexes currently employed.^4^ The use of somewhat larger iron oxide nanoparticles as T2 contrast agents is well established, with examples of such particles clinically approved as T2 contrast agents.^3^ Synthetic biology has also offered schemes for *in vivo* molecular imaging with MRI, including imaging tumors via selective expression of MRI contrast reporter genes or by modulating endogenous ferritin expression.^5–8^ In another study, expression of genetically encoded reporters based on gas vesicles for hyperpolarized xenon magnetic resonance imaging was shown in living cells with potential use for *in vivo* imaging via systemic injection of tumor-targeted encoded viral vectors.^9^ Nanomaterial synthesis and synthetic biology intersect with the intriguing concept of harnessing magnetotactic bacteria (MTB), a group of prokaryotes known for their ability to biomineralize pristine intracellular nanocrystals of magnetite (Fe_3_O_4_) with sizes ranging from 35-120 nm, as a biogenic source of high quality T2 contrast agents.^10–12^ These particles occur in chains and are surrounded by a phospholipid bilayer membrane, forming a structure called a magnetosome. Recent research demonstrated the display of targeting peptides on the surfaces of these magnetosomes, such as RGD and recently the GFR/HER2 targeting peptide P75.^13–16^ As targeting moieties, peptides offer several advantages, including their small size, high affinity, ease of modification, and low immunogenicity.^17–20^ Despite their promise, some peptides are challenging to attach to the surface of synthetic iron oxide nanoparticles while maintaining their structural confirmation, making transgenic expression on the surface of magnetosomes the most direct means to produce contrast agents in such cases. ^21,22^

This work provides the first report of magnetic nanoparticles displaying genetically-encoded pH low insertion peptide (pHLIP)^23^, a long peptide that is ill suited for chemical conjugation. Targeting mechanisms based on pH responsiveness hold particular promise as broadly tumor-selective MRI contrast agents, since they exploit the extracellular acidity associated with the tumor microenvironment for contrast enhancement.^24^ In separate genetically engineered magnetosomes, we incorporated the αv integrin-binding cyclic peptide iRGD, which is known to specifically target tumors by binding integrin-expressing cells in a neuropilin-1-dependent manner. Upon binding to αv integrins on the endothelium of tumor vessels, it is proteolytically cleaved within the tumor microenvironment, increasing affinity for neuropilin-1 and facilitating tissue penetration of co-administered or conjugated drugs or imaging agents.^25,26^ In addition to verifying the applicability of this technique for functionalization of magnetosomes with widely different peptide ligands, this approach provided a means to compare the performance of pHLIP against a peptide with known tumor targeting and penetration capabilities. We found increased binding affinity of both pHLIP- and iRGD-functionalized magnetosomes to cancer cells in low pH culture conditions, or to cancer cells expressing αv integrins, respectively, relative to non-functionalized magnetosomes. Further, this selectivity was also observed *in vivo* during experiments in which functionalized magnetosomes accumulated at tumor sites and produced measurable local decreases in T2 relaxation times. Because genetic engineering allows different peptides to simultaneously be displayed on individual magnetosomes, our findings suggest that pHLIP and iRGD could serve as useful components of a tailored peptide combination on magnetosomes acting as generalized tumor-selective T2 contrast agents.

## Results and Discussion

### Purified magnetosomes are suitable T2-relaxation contrast agents

To date, a range of magnetotactic bacteria strains have been discovered that all appear to synthesize magnetosomes. In this work, we employed *Magnetospirillum magneticum* AMB-1 cells, for which the complete genome sequence is available.^27^ Before undertaking the genetic modification of these bacteria to tag their magnetosomes with custom peptides in order to improve their targeting as tumor contrast agents, the structure and magnetic properties of AMB-1 and their extracted and purified magnetosomes were investigated. Using cryo-transmission electron microscopy (cryo-TEM), AMB-1 were imaged prior to magnetosome extraction to verify the formation of magnetosome chains which were then analyzed in more detail after harvesting and purification (see Materials and Methods). We observed 200-500 nm long chains of individual iron oxide crystals, encompassed by a 5–6 nm protein-rich phospholipid bilayer that made up the magnetosome membrane (**Fig. 1a**). The micrographs of the magnetosomes purified in this study showed individual grain sizes on the order of 40 nm, a size consistent with the range encountered in wild-type MTB and suggestive of their function as carriers of high-stability natural remanence. At physiological temperatures and long timescales, the critical size above which individual grains of cubic magnetite are no longer superparamagnetic is 25–30 nm,^28^ and pseudo single domain and multidomain behavior is expected at lengths above approximately 80 nm.^29^ The size of the isolated particles falls between these values, suggesting they should exist in a stable single domain (SSD) state. This characterization was corroborated by observations of hysteresis at room temperature when vibrating sample magnetometry (VSM) was performed on samples prepared in gels to prevent their physical rotation (**Fig. 1b**). Because individual iron oxide crystals within a magnetosome chain are only separated by a few nm, magnetostatic interactions have been observed to affect the orientation of the moment of each crystal, resulting in alignment along the axis of the chain.^30^ The same magnetic properties were retained for the purified magnetosomes, indicating that they maintained their structural integrity (**Fig. 1b**).

**Figure 1.**
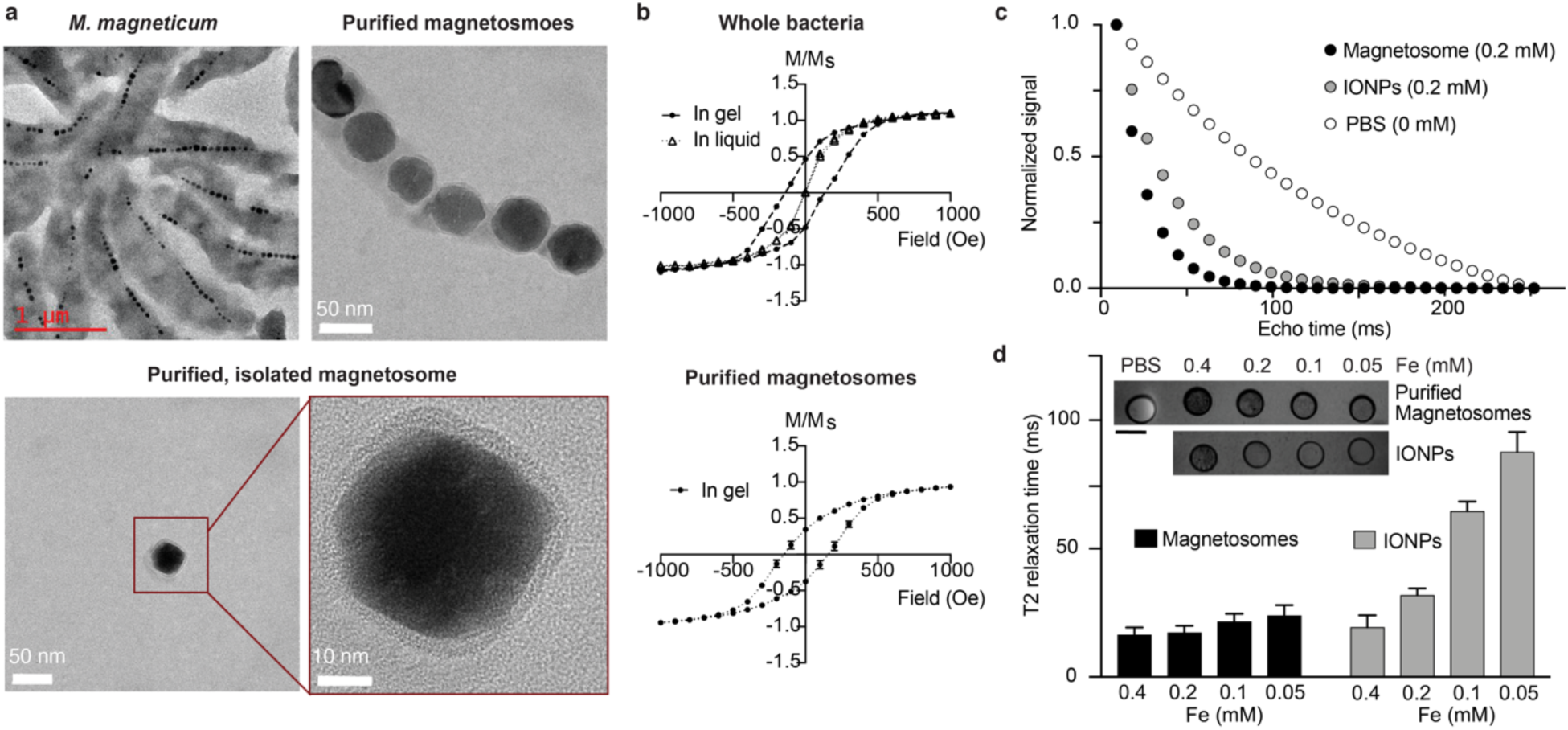
Magnetosomes are suitable T2 - weighted contrast agents. **a)** Transmission electron micrographs of *M. magneticum* before (left) and after (right) purification, showing an individual magnetosome. A zoomed-in TEM image of an isolated magnetosome core with lipid bilayer is shown in the bottom. **b)** The magnetic properties of intact AMB-1 cells (top) and purified magnetosomes (bottom) were characterized by vibrating sample magnetometry (VSM). Intact cells (top) were first measured in liquid and then immobilized in a gel to prevent physical rotation, allowing observation of their hysteresis (n=3 samples of one culture). Purified magnetosomes showed similar behavior, indicating that the structural integrity of the magnetosomes was maintained. **c)** The determined magnetic properties suggested their use as T2-weighted contrast agents which was assessed in in T2-relaxation time measurements and compared with samples of individually suspended IONPs with similar iron oxide nanoparticle core size (25nm). Samples were immobilized in tubes containing agar and the shortening of T2 relaxation time (decay of signal amplitude to 1/e) was determined, yielding 17.3±2.6 ms for the magnetosomes, compared to 31.9±2.6 ms for 25 nm large iron oxide nanoparticles, for both samples at 0.2 mM Fe content. The mean is derived from the average T2 relaxation time per pixel in the measured region of interest. **d)** T2 relaxation time increases, and thus, the effect on T2 relaxation time reduction decreases, with decreasing iron concentration, as to be expected. While the commercial IONPs show a fairly linear increase with decreasing iron concentration, a muted decrease was observed for purified magnetosomes, which might be attributed to local clustering effects. The inset (top) shows the T2 intensity of cross-sectional scans of the tubes for the respective concentrations of the two samples.

The magnetic properties of magnetosomes have prompted previous investigations into their suitability as contrast agents, both for magnetic particle imaging (MPI) and MRI.^11^ In a previous study, magnetosome contrast enhancement was determined to be slightly elevated compared to the former iron oxide-based “gold-standard”, Resovist, with a hydrodynamic diameter between 45 and 60 nm and an iron oxide nanoparticle core between only 5 and 6 nm.^11,31^ We measured an increased reduction of T2-weighted signal relaxation time of 17.3±2.6 ms for purified magnetosomes with an iron oxide core diameter of approximately 40 nm at an iron concentration of 0.2 mM. This effect on T2 relaxation time reduction exceeded that of commercial, well dispersed magnetite particles with a similar core diameter of 25 nm, for which a value of 31.9±2.6 ms was measured at the same iron concentration (**Fig. 1c,d**). This result is consistent with the capacity of purified magnetosomes to mediate a T2 relaxation time reduction and their suitability as a core material for functional MRI contrast agents.

### Engineered peptide-displaying magnetosomes retain wild-type magnetic properties

After characterizing the properties of the native AMB-1 magnetosomes, we sought to genetically engineer AMB-1 bacteria to produce peptide-functionalized magnetosomes. The technique to display peptides on the surface of magnetosomes has been well established during the past decade.^32,33^ Several studies have suggested that a magnetosome-specific protein called mamC is the most abundant protein on the magnetosome surface^34^, and thus it has been used previously for display. As a preliminary validation, we sought to display a his-tag on the magnetosome. To do this, we fused the his-tag sequence to the C-terminal of the mamC gene immediately before the stop codon, and placed this tag under the regulation of the IPTG-inducible ptac promoter. The resulting plasmid was then conjugated into *Magnetospirillum magneticum* AMB-1. The proteins from the purified magnetosomes were collected and analyzed by SDS-PAGE and Western blotting (**Fig 2a**). These results indicate that our experimental protocols indeed fuse peptides to the magnetosomes.

**Figure 2.**
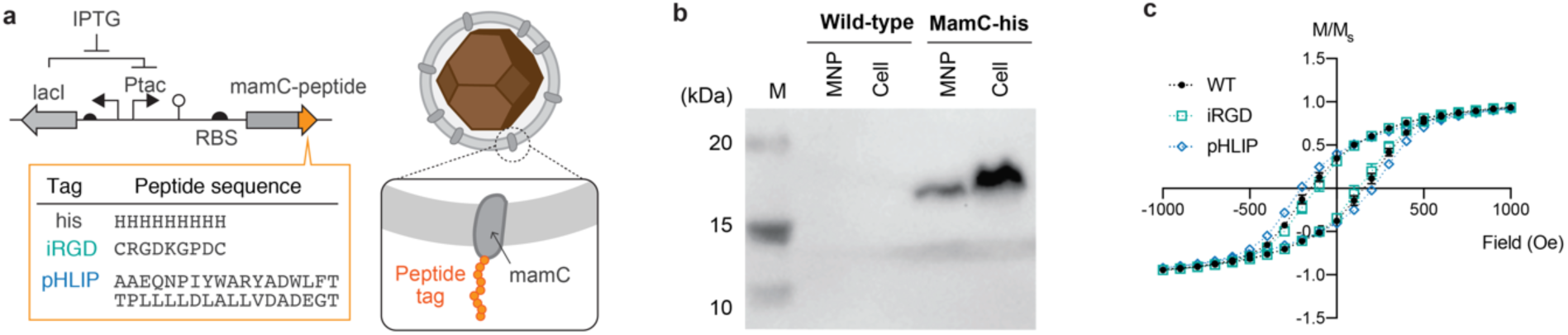
Genetic engineering of AMB-1. **a)** Peptide tag sequence (his-tag, iRGD, and pHLIP) was fused to the C-terminus of the mamC gene and placed under an IPTG-inducible P_tac_ promoter. **b)** Western blot with anti-his antibody of magnetosome nanoparticle binding proteins (“MNP”) or cell lysates (“Cell”) extracted from wild-type or MamC-his expressing strain. **c)** The magnetic properties of peptide-fused magnetosome were assessed and compared to the previous measurements of the hysteresis of wild-type magnetosomes. The behavior is largely consistent, suggesting that the genetic alterations did not affect magnetosome synthesis by the bacterial cells.

We proceeded to design DNA sequences to display pHLIP and iRGD on magnetosomes to add relevant functionality and selectivity in context of tumor targeting contrast agents, both for acidic tumor tissues and integrin-expressing cancer cells, respectively.^25,35–37^ pHLIP and iRGD peptide-encoding DNA sequences were generated by backtranslating using a GC rich codon to optimize expression in *M. magneticum*. These DNA sequences were fused to the C-terminus of the mamC gene immediately before the stop codon and after a linker sequence (**Fig. 2b**). Except for the peptide sequence, the construct is identical to the one validated for the his-tagged magnetosomes. The pHLIP- and iRGD-encoding plasmids were conjugated into *M. magneticum*, and magnetosomes from the cells were collected, purified, and thoroughly washed. To verify that genetic modification of their surface proteins did not alter properties relevant to their function as T2 contrast agents, we acquired hysteresis curves for these genetically engineered pHLIP- and iRGD-magnetosomes, which yielded comparable results to the wild-type magnetosomes (**Fig. 2c**). The saturation magnetization, as determined by these measurements and quantification of iron content via inductively-coupled plasma mass spectrometry (ICP-MS), suggested values of 111 emu/g_Fe_ for wild-type magnetosomes, 91.2 emu/g_Fe_ for iRGD-expressing magnetosomes, and 96.3 emu/g_Fe_ for pHLIP-expressing magnetosomes. These values indicate that the high saturation magnetization values of wild-type MTB magnetosomes are retained upon expression of iRGD or pHLIP and are consistent with the majority of iron in the particles occurring as biomineralized magnetite (115 emu/g_Fe_ in bulk).^38^

### Peptide-fused magnetosomes serve as functional binding agents

Next, we tested whether our selected peptides, pHLIP and iRGD, retained their functionality when displayed on the surface of purified magnetosomes. The unique characteristic of pHLIP is its ability to associate with lipid bilayers as an unstructured monomer at neutral pH, while inserting across a bilayer or membrane as an alpha-helix in acidic conditions. As a result, magnetosomes displaying pHLIP are expected to show a pH-dependent fusion to cell membranes compared to unmodified magnetosomes. Thus, we cultured cells of the human breast cancer cell line MDA-MB-231, both at a standard pH of 7.5 and at a low, slightly acidic pH of 6.5, which is representative of average tumor acidity.^39^ We fluorescently labeled magnetosomes with and without expression of pHLIP using a lipophilic membrane dye (DiI) inserted into the membrane of the magnetosome. We incubated both wild-type and pHLIP-functionalized magnetosomes with MDA-MB-231 cells at low pH and standard pH and analyzed the binding efficiency by flow cytometry. Our data revealed a significant increase in bound magnetosomes to cells for pHLIP-functionalized magnetosomes at low pH compared to neutral pH, as well as compared to pure magnetosomes at low pH, demonstrating pH-activated binding to cancer cells *in vitro* (**Fig. 3a**). As expected, there was no observable difference in the binding efficiency of pure magnetosomes with cancer cells between these two different pH levels.

**Figure 3.**
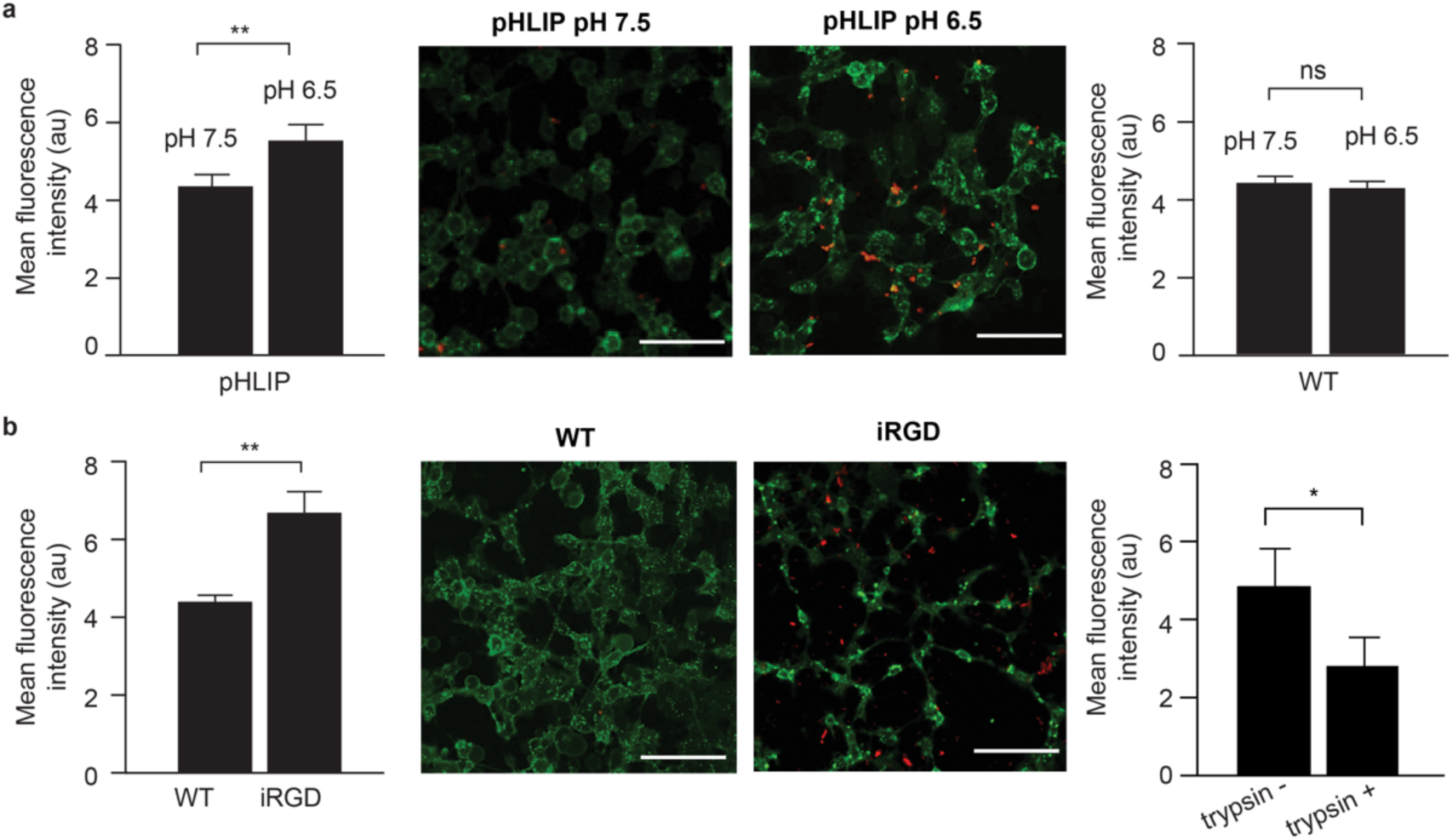
Magnetosomes functionalized with tumor-targeting peptides pHLIP and iRGD show increased binding affinity to cancer cells. **a)** pHLIP and wild-type magnetosomes were fluorescently labeled with a membrane dye (green) and incubated with MDA-MB-231 breast cancer cells at standard (pH 7.5) and low pH (pH 6.5). Increased binding for pHLIP magnetosomes at low pH was measured by flow cytometry, compared to their binding at standard pH (left, p = 0.0078, unpaired, two sample t test). This can be also seen in fluorescence images of cells (green) with magnetosomes (red) shown on the right (scale bars = 100 µm). No such pH dependency was observed for wild-type magnetosome (right, p = 0.6420, unpaired, two sample t test) **b)** In another experiment, iRGD magnetosomes were incubated with the same cell line, known to express αv integrins. A significantly increased selectivity to these cells was observed for iRGD magnetosomes compared to wild-type magnetosomes left, p = 0.0033, unpaired, two sample t test), which is also reflected in the microscopy images on the right (scale bars = 100 µm). Binding efficiency significantly dropped when iRGD-functionalized magnetosomes were incubated with trypsinized cells, and thus bearing cleaved integrins, compared to untrypsinized cells with preserved integrin expression (right, p = 0.0411, unpaired, two sample t test).

Analogously, we assessed the functionality of iRGD displayed on magnetosomes by testing the specificity of binding to αvβ3 expressing cells, such as the MDA-MB-231 cells used in the previous experiments.^40,41^ A 1.53-fold increase in binding efficiency was measured by flow cytometry for iRGD-functionalized magnetosomes compared to wild-type magnetosomes (**Fig. 3b**). Analysis of fixed samples by confocal imaging also revealed the localization of the magnetosomes at the cell surface. Further, trypsin treatment, which has been shown to cleave cell-surface integrins non-specifically,^42^ resulted in a 42% decrease in the fluorescent intensity of iRGD-functionalized magnetosome binding, further supporting our hypothesis of integrin-selective targeting of iRGD-functionalized magnetosomes (**Fig. 3b**).

### Magnetosomes fused with tumor-targeting peptides serve as potential *in vivo* cancer imaging agents

Finally, we tested peptide-functionalized and non-decorated magnetosomes *in vivo* by intravenously injecting suspensions of the respective purified and near infrared (NIR) fluorescently-labeled magnetosomes into mice bearing flank xenografts of the human ductal carcinoma (mammary gland origin) cell line MDA-MB-435S. These cells strongly express the integrin αvβ3,^43,44^ and thus, we expected to observe an increase in tumoral accumulation of the iRGD-fused magnetosomes relative to non-decorated magnetosomes. To assess biodistribution, organs were harvested 6 and 24 h post injection and analyzed with a NIR scanner (**Fig. 4a,b**). Both pHLIP and iRGD peptide-functionalized magnetosomes showed a trend towards increased accumulation at tumor sites compared to pure magnetosomes; specifically, approximately 1.5-fold and 2-fold increases in tumoral accumulation for pHLIP and iRGD, respectively, were measured after 6h (**Fig. 4c**). This data is consistent with values found in comparable previous studies, including a large literature survey on tumor accumulation of nanoparticles that broadly suggest a 1.5- to 2-fold increase in tumor delivery for targeted versus non-specific nanoparticles.^17,45^ A decrease in signal after 24h suggested rapid clearance of the magnetosomes.

**Figure 4.**
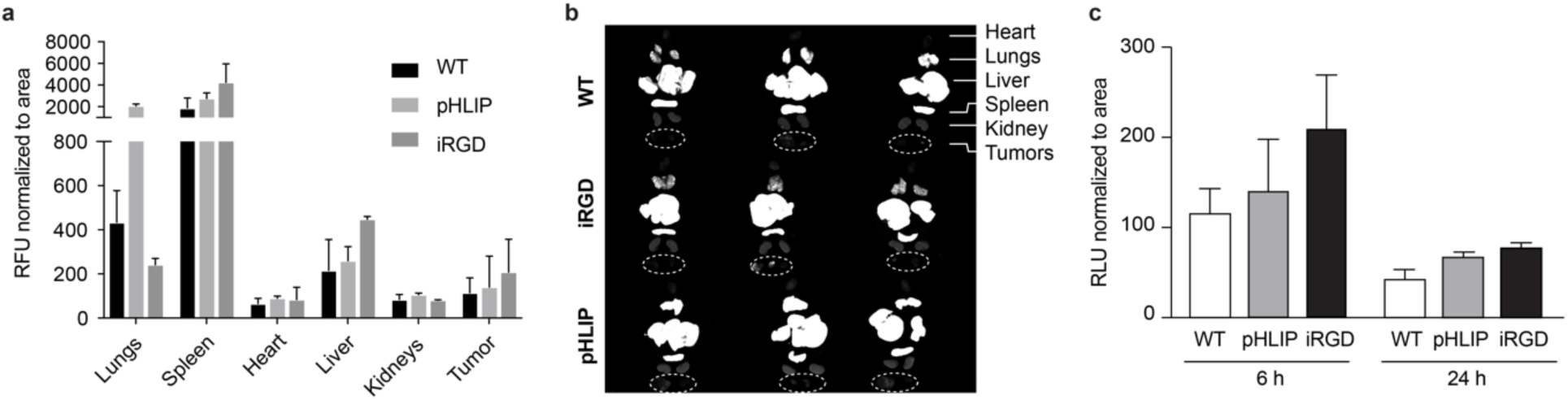
Peptide-fused magnetosomes show increased accumulation in tumors. **a)** Near infrared (NIR)-labeled (lipophilic dye Cy7.5) magnetosomes were intravenously injected into mice (n=3 each group) bearing MDA-MB-435S flank tumors. Organs were harvested after 6h and measured with a NIR scanner, with relative fluorescence units (RFU) normalized to the area of each organ. **b**) An intensity plot is shown for a scan at 800 nm emission wavelength on the harvested organs. Tumors are circled in white. **c**) Comparison of tumoral accumulation at 6h and 24h post injection. An increased accumulation of peptide displaying magnetosomes, although not statistically significant, was found in tumors compared to wildtype magnetosomes, with levels diminishing rapidly within 24h (n=3 each group).

Based on these results, we decided to analyze the tumors within the first 6 h post injection with T2-weighted image analysis in a 7T whole mouse MRI scanner. A darkening of the tumor regions was observed for all magnetosome types, as shown in scans in **Figure 5a**, while the strongest enhancement was observed for iRGD-fused magnetosomes, in line with the results of the biodistribution analysis and similar to results obtained *in vitro* for the αvβ3 expressing MDA-MB-231 cell line. We then quantified the impact on T2-relaxation time reduction, i.e. the extent to which iRGD-functionalized magnetosomes darken the targeted tumor region (**Fig. 5b**). Compared to wild-type magnetosomes, the decrease in relaxation time increased by 2.02-fold, although not with statistical significance, with three animals in each group bearing two flank tumors.

**Figure 5.**
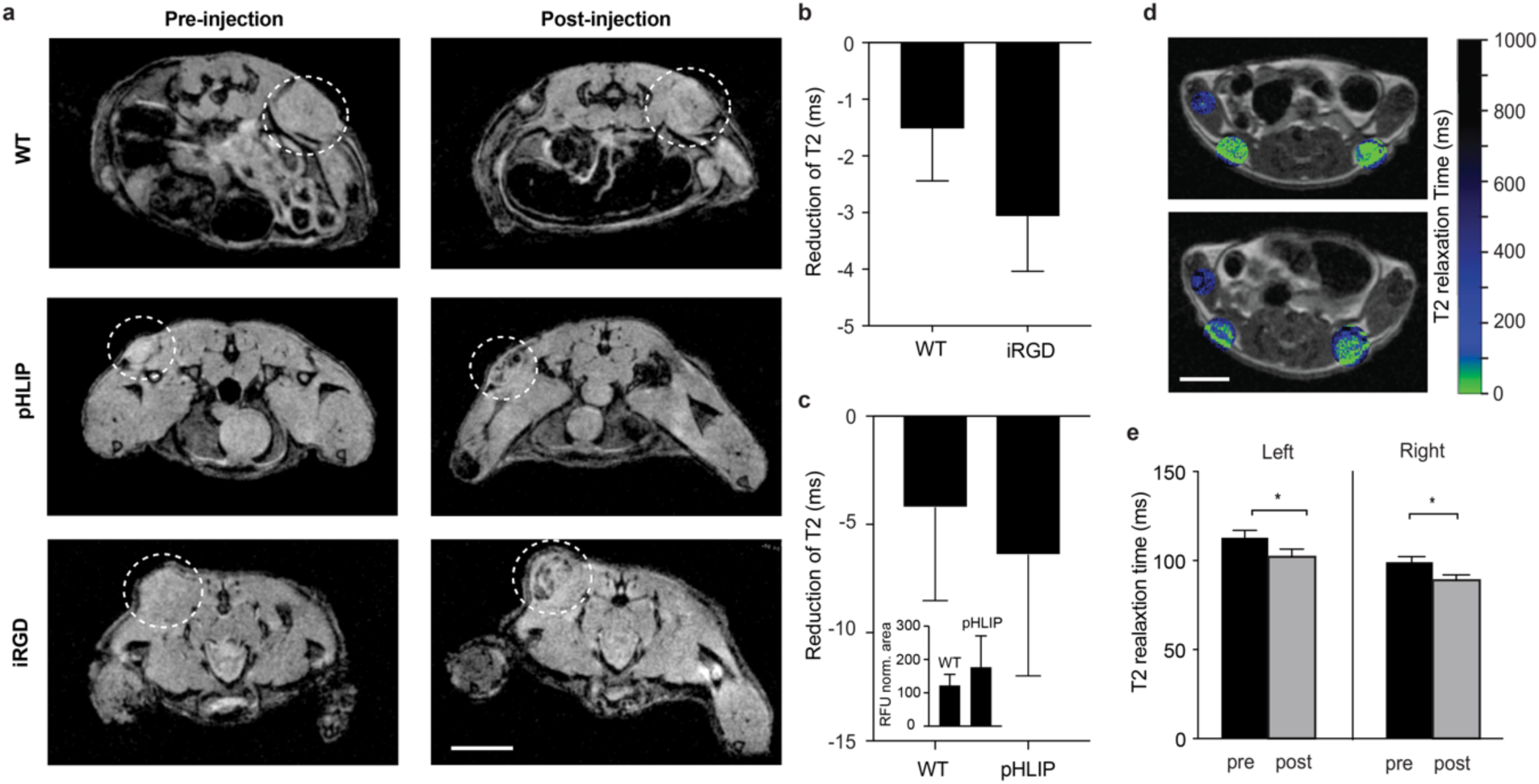
Peptide-fused magnetosomes show enhanced T2 weighted contrast in two different tumor models. **a)** Mice bearing MDA-MB-453S flank tumors were imaged in a 7T MRI scanner and T2-weighted relaxation time was determined before and after intravenous injection of magnetosomes. An enhanced darkening, attributable to a decrease in T2 relaxation time can be observed for peptide-displaying magnetosomes relative to wild-type magnetosomes (scale bar = 10 mm, same scale for all images, representative scans are shown for each group, n = 3 each group). **b)** A 2.02-fold increase of the reduction of T2 relaxation was found across tumors in mice that were injected with iRGD magnetosomes compared to those exposed to wild-type magnetosomes (p=0.277, not significant according to unpaired, two-tailed student’s t-test, n=3 animals, each, two flank tumors per animal). **c)** pHLIP functionalized magnetosomes were injected into mice bearing LS174t flank tumors and an increased tumor accumulation of pHLIP functionalized over non-functionalized magnetosomes 6h post injection was observed (inset). Overall a T2 relaxation time reduction of 7.72% (6.40 ms) was measured (p= 0.309, not significant according to unpaired two tailed student’s t-test, n=3 for the control and n=4 for the experimental group), demonstrating the potential of pHLIP displaying magnetosomes for T2 weighted *in vivo* tumor imaging. **d)** The magnitude of T2 relaxation time reduction was quantified and mapped in this example onto scans acquired by fast spin-echo multi-slice scan sequence (fsems) of a mouse with two flank tumors, injected with pHLIP magnetosomes (scale bar = 10 mm, same scale for both images). The T2 relaxation time before and 5h after injection is shown in **e**).

To demonstrate the impact of low pH targeting *in vivo*, we characterized the performance of the pHLIP-functionalized magnetosomes in a flank tumor model of the LS174T colorectal cancer cell line, which only faintly expresses the αv integrin.^46^ To further increase the potential contrast enhancement, we concentrated our magnetosome formation from 0.5 mg/ml to 1.5 mg/ml compared to the previous trial and injected 0.3 mg per mouse. An increased accumulation at tumor sites was again measured for pHLIP-functionalized magnetosomes over non-functionalized magnetosomes (**Fig. 5c**, inset). The T2-relaxation was reduced for pHLIP magnetosomes by 7.72% (6.40 ms), however, not significantly across the small number of animals (n=3 for the control and n=4 for the experimental group, **Fig. 5c**). An example scan of two flank tumors is shown in **Figure 5d** with an overlaid T2 relaxation time map, which was quantified across 6 slices (**Fig. 5e**). Follow up studies with larger cohorts of animals could be suggested for higher powered statistics; however, these results already indicate the potential of peptide functionalized magnetosomes as T2-weighted contrast agents for *in vivo* tumor imaging.

## Conclusion

We have demonstrated the design, synthesis, and application of T2-weighted tumor imaging probes produced through genetic manipulation of MTB to display the tumor-targeting peptides pHLIP or iRGD on their magnetosome membranes. This genetic modification was found to leave the magnetic properties of these structures substantially unaltered. The intended functionality of both peptides was verified experimentally *in vitro.* Animal studies employing small numbers of mice also showed indications of tumor selectivity and enhanced T2-weighted contrast *in vivo*. The up to 2-fold enhancement in accumulation for targeted magnetosomes is consistent with the range of delivery enhancement previously suggested in literature for tumor targeting nanoparticles.

This approach combines the biomineralized cores of magnetosomes, along with their uniquely advantageous magnetic properties, with genetically engineered surface functionalizations that impart targeting and other useful capabilities. The versatility and flexibility to specifically tailor and rapidly synthesize new peptide-functionalized imaging probes with such pristine magnetic properties has great potential in clinical applications as targeted diagnostic agents, specifically when incorporating targeting peptides, that are complex to synthesize synthetically. The simultaneous incorporation of multiple peptides to obtain multiplexed functionality could open new avenues in multimodal diagnostics.

## Materials and Methods

### Bacterial media composition

Modified Magnetosome Growth (MMG) media was prepared by a slight modification of Magnetospirillum Growth Media (MSGM).^47^ First, MMG was prepared by adding the following to 1L distilled water: 0.68 g KH_2_PO_4_ (Fischer), 0.34 g NaNO_3_ (Fischer), 0.37 g tartaric acid (Alfa-Aesar), 0.37 g succinic acid (Sigma-Aldrich), 0.05 g sodium acetate (Sigma-Aldrich), 5 mL ATCC Trace Mineral Solution. The pH was then adjusted to 6.8 by 10 N NaOH, and the solution was autoclaved at 121°C for 40 min. The MMG could be kept on the shelf for ∼2 months. Just before each bacterial culture, ATCC Vitamin Supplement (ATCC MD-VS, 100 to 1000x dilution), 10 mM ferric quinate (100x dilution, final concentration of 100 µM), ascorbic acid (see “Bacterial strains and culture conditions” for the concentration), and appropriate antibiotics or inducers were freshly added to the MMG.

Agar media with final concentration of 0.8% agar was prepared by adding 8 g of Bacto Agar (Difco) to 1 L MMG-base and autoclaving at 121°C for 40 min. After the media was cooled below 60°C, ATCC Vitamin solution (1000x), 10 mM ferric quinate (100x), and appropriate antibiotics were added and poured into a plastic dish. (Note that ascorbic acid is not included). The plates were stored at 4°C, and warmed to 30°C just before plating.

Stock solutions were prepared as follows: 10 mM ferric quinate (100x) was prepared by adding 0.27 g of FeCl_3_ 6H_2_O and 0.19 g of D-(-)-quinic acid, sterile filtered, and used within 2 weeks. 1000x ascorbic acid solution (1 g/L for seed culture or 20 g/L for passage culture) was prepared and sterile filtered just before each culture. 100× 2,6-diaminopimelic acid (DAP) was prepared by adding 0.285 g of DAP to 100 mL distilled water while heating, followed by sterile filtration. Antibiotics used (in their final concentration) were 5 µg/mL Kanamycin and 5 µg/mL Gentamycin for liquid culture, or 10 µg/mL Kanamycin and 10 µg/mL Gentamycin for agar culture.

### Bacterial strains and culture condition

*E. coli* NEB 10-beta (NEB) was used for cloning, and WM3064 (also known as BW29427) (thrB1004 pro thi rpsL hsdS lacZΔM15 RP4-1360 Δ(araBAD)567 ΔdapA1341::[erm pir], derived from W. Metcalf)^47^ was used for plasmid conjugation. WM3064 harbors RP4 conjugation machinery in the genome, and lacks dapA gene and therefore requires DAP to grow.

*M. magneticum* AMB-1 was purchased from ATCC and cultures were started by streaking glycerol stocks on MMG agar or inoculating glycerol stocks directly into a liquid culture. After streaking the cells, the plates were inserted in campy pouch (BD) and incubated at 30°C for 5-7 days to obtain single colonies. Tiny colonies are visible after 4-5 days, and growing them for additional 2-3 days (total 7 days) resulted in fully-grown colonies. The plates were pulled out from the bag for further experiments. The plates were kept in a bag flushed with nitrogen to keep a microaerobic environment at room temperature and used within 1 week.

For the seed culture in liquid media, MMG media was prepared just before inoculation by adding the following to MMG-base (described in section “Bacterial media composition”): ATCC Vitamin solution (1000x), 0.1 g/L ascorbic acid (1000x), 10 mM ferric quinate (100x) and appropriate antibiotics or inducers. Seed culture was performed in 5 mL of MMG media using 10 mL gas vials. Single colonies were inoculated into the media, and the rubber-cap was tightened. The headspace was replaced with N2 by vacuuming the headspace and adding N2 gas, then repeated three times. 250 µL of air (approximately 50 µL of O2, which is 1% of the headspace) was injected into the headspace using a needle and a syringe. The vial was placed into a water bath at 30°C and shook at 60 rpm. This seed culture was incubated for 3 days.

Second-passage cultures were performed in either 5 mL or 100 mL culture. MMG media was prepared with slight modification, by adding 1000× 2 g/L ascorbic acid (final concentration 20 mg/L). For 100 mL culture, 100 mL MMG media using a 170 mL glass jar, sealed by rubber cap (Sigma-Aldrich). The cells were inoculated by 1:50 dilution (by adding 2 mL seed culture), the headspace was replaced with N2, 700 µL O2 was added, and cultures were incubated at 30 °C.

For large-scale culture (third-passage culture), 2.4 L MMG media was prepared in a 2.8 L flask. Fifty mL of second-passage culture was added, the flask was sealed with a rubber stopper, and the headspace was replaced with 1% O_2_ and 99% N_2_ using a needle through the rubber stopper. The culture was cultured at 30°C, 60 rpm.

### Magnetosome purification

Magnetosomes were purified from *M. magneticum* based on the method reported by Borg et al.^34^ with modifications. Four batches of *M. magneticum* culture were collected in 1 L bottles with centrifuge at 5000g for 20 min. The pellet from 1 L culture was resuspended with 60 mL of 20 mM HEPES (pH 7.4) with 1 mM EDTA (HEPES-E buffer), and 30 mL of culture was dispensed in two 50 mL tubes (or 60 mL culture was collected into 250 mL bottles), and centrifuged at 4700g for 20 min. The cell pellet was kept at −80°C if not immediately used.

The cell pellet was resuspended in HEPES-E supplemented with 0.1 mM PMSF by vortexing. The resuspension was passed through a French Press at 20 kpsi. The lysate was transferred to 250 mL bottles, and was sonicated on ice for 1 min at 10% amplitude with 1-sec ON/OFF interval. Lysates were transferred to 50-mL falcon tubes and placed next to a strong magnet overnight at 4°C. The supernatant was aspirated, and the magnetic fraction was resuspended into 10-mL HEPES-E. The solution was sonicated on ice for 30 sec at 10% amplitude with 1-sec ON/OFF interval. This was placed next to a strong magnet for 4 hours at 4°C, and this washing step was repeated 3-10 times. After washing, the magnetic fraction was resuspended a final time in distilled water or PBS and kept at 4°C until use.

### Plasmid construction

Plasmids and genetic construct used in this study are described in Supplementary Table 1. pMGA vector was constructed by merging the pMGT vector^48^ and p15A plasmid. The *mamC* gene was PCR-amplified from the *M. magneticum* genome and inserted under the IPTG-inducible Ptac/LacI promoter. The peptide sequence for the his-tag, iRGD and pHLIP were (in single-letter amino acid abbreviations) HHHHHHHHH, CRGDKGPDC^26^ and AAEQNPIYWARYADWLFTTPLLLLDLALLVDADEGT^35^, respectively. The DNA sequence for these peptide tags were generated by backtranslating using the GC-rich *Streptomyces coelicolor* codon usage table. The pHLIP-tag were fused right after the C-terminal of *mamC* CDS, while the his-tag and iRGD-tag were fused to *mamC* after an AGGS peptide linker. The DNA sequence of the fused *mamC*-peptide CDS are shown in Supplementary Table 2.

### Cell culture

MDA-MB-231 and MDA-MB-435S cells were cultured in high glucose Dulbecco’s Modified Eagle’s Medium (DMEM, Gibco) supplemented with 10% foetal bovine serum (FBS) in a humidified atmosphere with 5% CO_2_ at 37°C. MDA-MB-435S cell cultures were supplemented with 1% penicillin-streptomycin (CellGro). LS174T were cultured in Eagle’s Minimal Essential Medium (ATCC) supplemented with 10% FBS (Gibco) and 1% penicillin-streptomycin (CellGro). All cells were passaged at 80% confluency.

### Measurement of magnetic properties of MTB and purified magnetosomes

A Microsense vibrating sample magnetometer (EZ VSM) measured moment versus field curves at room temperature for 100 µL samples of MTB and magnetosomes prepared in modified NMR tubes. Samples were affixed with silicone vacuum grease to a vibrating sample rod shortened to ensure centering in the gap of the electromagnet between the sensing coils. Where indicated, to prevent physical rotation of MTB or magnetosomes, samples were prepared in 2 to 4 % agarose gel by adding agarose powder to the samples, briefly microwaving them (30 to 60s) while held upright in a thin layer of water in a glass petri dish, and placing them on ice to gel. Diamagnetic signal was subtracted by linear fits of the data at high field magnitudes, and M/Ms curves were obtained by normalizing to the saturation moment indicated by the intercepts of these linear fits.

### *In vitro* evaluation of magnetosome targeting

MDA-MB-231 cells were cultured to 80% confluency on coverslips in 6-well plates and stained with 200µL complete DMEM containing 1µL DiO at a concentration of 1 mM (Sigma) to visualise the cell membrane. After 45 min incubation in a humidified atmosphere with 5% CO2 at 37°C, the cells were washed twice with 1X PBS. Functionalized magnetosomes were stained with DiI (Sigma), incubated at room temperature with gentle agitation for 30 min and were washed twice with PBS. For iRGD-fused magnetosomes, cells were treated with 400 µL of magnetosomes in 1 mL complete DMEM on ice. For pHLIP-fused magnetosomes, cells were treated with 50 µL of magnetosomes in 1 mL complete DMEM on ice at pH 6.5 or 7.4. The pH 6.5 medium was prepared by adding 20 mM HEPES and 20 mM MES to complete DMEM. Cells treated with unfunctionalized magnetosomes were used as negative controls. After one hour incubation, the magnetosome solution was removed and cells were washed twice with PBS. The cells were then fixed with 4% paraformaldehyde (PFA) on ice for 5 min and then washed twice with PBS. The coverslips were mounted on glass slides and were viewed under a spinning disk confocal microscope.

### Quantification of magnetosome targeting by Flow cytometry

For quantitative analysis of magnetosome binding, MDA-MB 231 cells were cultured to approximately 80% confluency. Cells were detached and transferred to tubes before treatment with stained functionalized magnetosomes. For iRGD-fused magnetosomes, 125,000 cells were treated with 25 µL of magnetosomes in 500 µL complete DMEM on ice. For pHLIP-fused magnetosomes, 500,000 cells were treated with 25 µL of magnetosomes in 500 µL complete DMEM on ice at pH 6.5 or 7.4. Cells treated with unfunctionalized magnetosomes were used as negative controls. After 1 h incubation, the magnetosome solution was removed and cells were washed twice with PBS. The cells were then fixed with 4% PFA on ice for 5 min, washed twice with PBS and resuspended 2% bovine serum albumin (BSA) in PBS. Cells were analyzed using a 561 nm excitation laser and 586/15 filter on a BD LSRFortessa. Untreated cells were used to set the gate on live cells (FSC/SSC). The fluorescence emission of 10,000 cells was recorded and targeting data are reported as mean fluorescence intensity (MFI). For all samples and controls Student’s t-tests were performed.

### ICP-MS

Samples of 50 µl were prepared in glass vials and evaporated overnight in a ventilated oven or on a hot plate. 200 µl of concentrated HCl (37% w/w) was added to dissolve the iron and diluted in a solution of 2% HNO3 in purified water. Analysis was carried out using ICP-TOFMS instrument (icpTOF, TOFWERK AG, Switzerland) or ICP-MS (Agilent 7900 ICP-MS). A reference solution was used at the beginning, middle, and end of sample measurements as a quality control. For calibration, five reference solutions containing different concentrations of Fe, and Rh, Co or In as internal standards were prepared.

### *In vivo* cancer model studies

Female nude mice (4–6 weeks, Taconic) were inoculated bilaterally with 3 × 10^6^ MDA-MB-435S or LS174T cells per flanks. Tumor growth was monitored and 10-14 days after inoculation, biodistribution and MRI studies were scheduled. Purified magnetosomes were labeled with a near infrared (NIR) lipophilic membrane dye (DiR’; DiIC18(7); 1,1’-Dioctadecyl-3,3,3’,3’-Tetramethylindotricarbocyanine Iodide, ThermoFisher) at a volume ratio of 1:2000, washed three times, and suspended in PBS. Iron concentrations were assessed by ICP-MS, and solutions were adjusted to yield 0.5 mg/ml Fe or 1.5 mg/ml Fe (two trials) respectively. Suspensions were administered intravenously via tail vein injections at a maximum volume of 200 µl and an iron mass of 0.1 mg or 0.3 mg.

### Magnetic resonance imaging

For MRI phantoms, serial dilutions of 0.4 mM Fe stock suspensions of magnetosomes and commercially available IONPs (SHP-25, Ocean Nanotech) were prepared in PBS. 200 µl of each sample was added into PCR tubes containing pre-weighted agar yielding final suspensions of 1 wt%. Samples were thoroughly mixed and sonicated, heated in a microwave, and cooled on ice to allow gelling. Imaging phantoms were prepared by inserting PCR tubes into 50 ml tubes, filling them with with 1 wt% agar, followed by heating in a microwave and gelling on ice. Phantom imaging experiments were then performed on a 9.4T small animal scanner (94/30 USR, Bruker, Ettlingen, Germany), equipped with a 87mm quadrature volume resonator. Multi echo multi slice sequence (MEMS) was used to evaluate T2 relaxation time, with the following parameters: repetition time TR = 2000 ms, number of echoes = 28, echo time TE = 9 ms to 252 ms with a 9 ms increment, data matrix = 320×120, Field of View (FOV) = 80×30 mm, 5 slices, slice thickness = 1.5 mm, 4 averages in 16 min.

MRI experiments were performed on a 7T MRI whole mouse MRI system (Varian 7T/210/ASR, Varian/Agilent), equipped with a 38mm mouse body coil. Multi echo multi slice sequence (MEMS) was used to evaluate T2 relaxation time, with the following parameters: repetition time TR = 1500 ms, number of echoes = 28, echo time TE = 9 ms to 252 ms with a 9 ms increment, data matrix=128×128, Field of View (FOV) = 50×50 mm. T2 weighted images were acquired with fast spin-echo multi-slice sequence (FSEMS) with the following parameters: TR = 2000ms, TE =2 4ms, ETL = 4, kzero= 2, 256×256 matrix, FOV = 50×50 mm^2^, interleaved number of slices=20 with no gap and slice thickness=0.5mm, number of averages = 2. Images were converted to DICOM format for viewing and analyzing purposes.

During imaging, mice were anesthetized by inhalation of 2.5% isoflurane and maintained on 2% isoflurane during data collection. Hot air was delivered throughout the imaging session to provide heat. Scans were collected with respiratory gating (PC-SAM version 6.26 by SA Instruments Inc., Stony Brook, New York) to avoid confounding noise due to chest movement.

## Statistics and Data Analysis

All statistical analyses were performed in GraphPad (Prism 8.0). Statistical significance and individual tests are described in figure legends.

## Acknowledgements

We thank Dr. M.-A. Augath and Dr. A. Schröter from the Rudin Laboratory at ETHZ for assistance with MRI scans of phantoms, Dr. H. Fleming (MIT) and Dr. M. Christiansen (ETHZ) for critical reading and editing of the manuscript, Dr. L. Gonzalez and Dr. P. R. Leduc (CMU) for the pMGT plasmid, and the Koch Biopolymers & Proteomics Core for assistance. We thank the Koch Institute Swanson Biotechnology Center for technical support, specifically S. Malstrom in the Koch Institute Animal Imaging and Preclinical Testing core. S.S. gratefully acknowledges the support by the Branco Weiss fellowship. The work from M.F. and C.A.V. was supported by the Institute for Collaborative Biotechnologies through contract W911NF-09-0001 and W911NF-19-2-0026 with the U.S. Army Research Office. A.P.S. thanks the NIH Molecular Biophysics Training Grant and the National Science Foundation Graduate Research Fellowship Program for support. S.N.B. is a Howard Hughes Medical Institute Investigator. This study was supported in part by a Koch Institute Support Grant P30-CA14051 from the National Cancer Institute (Swanson Biotechnology Center), a Core Center Grant P30-ES002109 from the National Institute of Environmental Health Sciences, the Ludwig Fund for Cancer Research and the Koch Institute Marble Center for Cancer Nanomedicine.

---

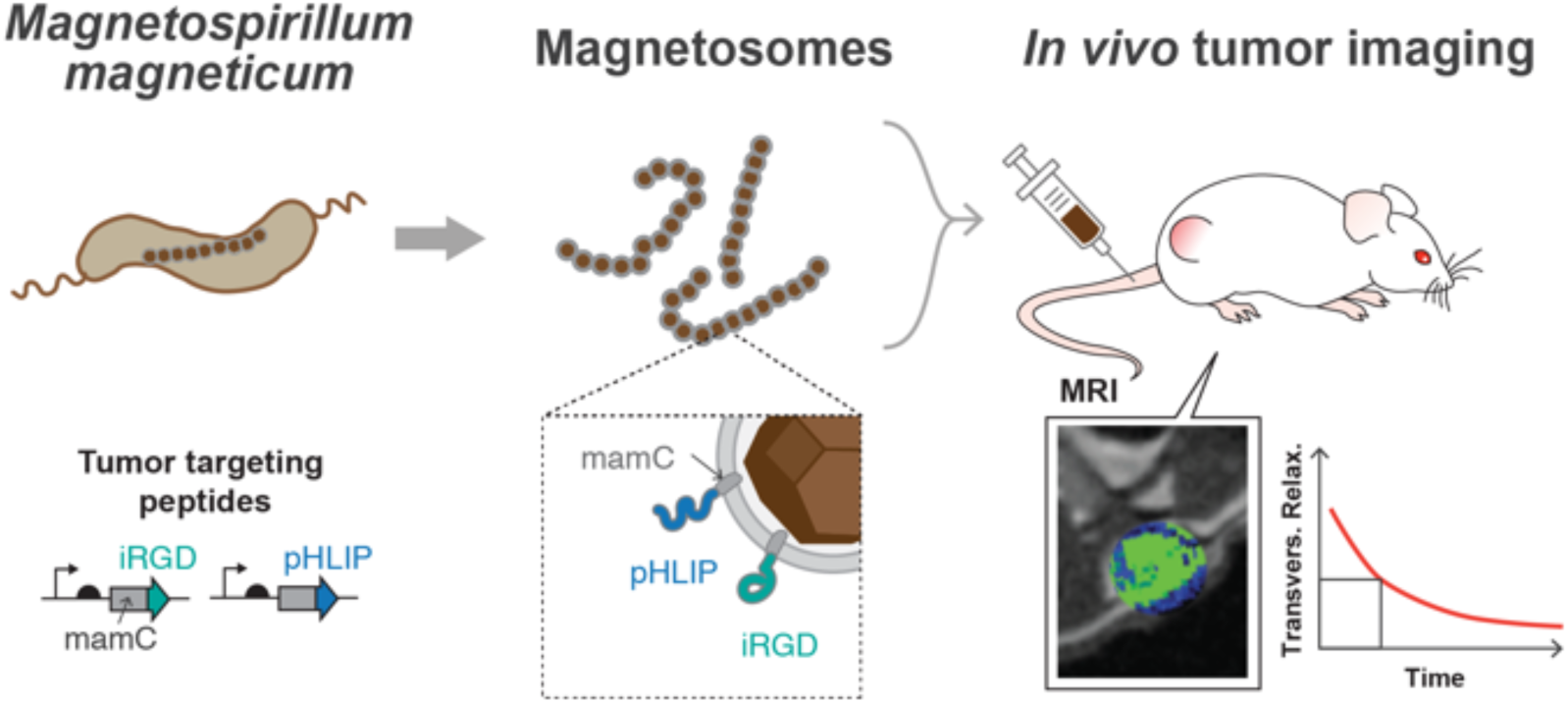
Graphical Abstract: Concept of genetically engineered magnetic nanoparticles with tumor targeting peptides as in vivo contrast agents.

