## Supplementary Table 1 for "Genetic encoding of targeted MRI contrast agents for *in vivo* tumor imaging"

**Supplementary Information**

**Supplementary Table 1. DNA sequence of the plasmid vector construct.**

| **Construct name** | **DNA Sequence** |
| --- | --- |
| pMGA vector | GGGTCCCCAATAATTACGATTTAAATTGGCGGGATCCAATCGCTCGAGCGCGAGGTCACAATTGGAGATGATCCGCCAGCGCCGAAGCCATCATAACTGCATTGAATAATTCAAAGTGCATTGAATAAGGCAGGGTGCATAAGGCGCAGCCTTTCGATTAACTCCATTTCCTGCGGCGACAACGTCGCCGCTTGCCCCCCCGACGGGGGTCAAGCGAGGCACAGGGACCCGCCGCCTCTTGGCGGTGGGTGGCCGCAGCTTGACGCCCGAACACTCCGCCGCCCAACCCTGCCTGCTCTCCTCTGTTCAATGAAATTGAACGGAGGAGATGCCTCCCGGCGAGGGCCTTTCTTTCTTGCATAACGTTCATGCTACGTTCCAACGGTTATACCGTATGAGCGTCGAACTGTATAATCGTTGAACAGTCGAACCATCCCCCCGCGCCAGCGCGGGGAGCAGGGATGGGTTCGGGGGCCGGAAGTTAGGCAAATCGCGGAGCCGTGGCGCCCTCGGTACATCCGCGAAATCGGTTGACACTTTGCCCCCAGAACGGCTGCAACCTACCGCTTGGCGACAGCCGCGTGACGGTACTTTTCCGCTGTGGATTCGGACACGAATATCTGTTCAGCAGCCAGGGCGCGGGCGATCTGGCGACGGCCGAGGCCCTGATGGGCCAATTCTGTGATCCGGTCTATGACCGCTTGGCTCATCGGCACCCGAGGACCGCCGATACCAGATTGGCGCACATCGCCCATGGTGGGTGGAACCGAGGGAGCCGGAGCCGGTGCAGATTTCTGCACAGTTGGTTCACGCCGTCGTTCTAGCGCATCGAGACGCTGCACGATCACGGCGAGCTGATCGTTGACGTATTGTCGCTCCTCGGACTGCGCAATTGTGATCAGAAATTCGAGGAGCTCGGCCACGGTGACGCCGCGTCGAAGGGCCATAGCCTGTATGTGGTCGCGGACCGCATCAGGACAGCGAATCGTCCATCTCGTGTCTGACACATTCCCGGTTTTTGTCTGACTCATCGTCGCCCCCAATGTCTGACAGATTCGAGAAAGAGTCATACCGGCTAACGCGAACGCTGACAACGAATGTCTGACAAAATGATGATTACGTCAGACATTGATGTCTGCCTAAATCGCAATAGTGTCTGACATGGTTTTTGGCGTTCCGTATATAAAATATGCGCCAAAATCTAGTGCGTCAGCCAAACGGCAAACGCAACCAGTTGACACCATTTCCGGTTTTATGAGACTGTGAAAAGACCAAAGGCCGCCTTGCGGCGGCCTAGGTGATTGTGGTCCCGGCGACCAAGGCTGCGAAAACTTGGCGGTCCAGCAGCAGAGGAAGCCCGTTTGAAAGGTCCGGCTTGCGCCGGAACCACCTAATGACAATAGCAATTCGCCGCACAGTTTTCAACACCCAGCCCGCCATGATTCGCGGCGGCGTTGAACGTTGAACAACAGACGTCTCAAACAGACACCCGCAAACGCGAACGGTATCGGTTGCAGGATGCAGCCGCTGAACTGCTGCCGGATTATCGGGTGCGGCATTGCCACCGCTCAATTGGATGTGGCTGTGGCGAGTTGGTCAGCGTTGTTACCAATGGGCAGTCGGCCAATTTCAAAGGCGTGCAGACCTGCGGCTCGGTCTGGCACTGCCCAATCTGTTCGGCAAAAATCAGCAATGTTCGGCGCGGCGAGCTGAACGAGCTGCTGCGCTGGGGGCGGGAGAGGGAGCTGATCCCGCTGATGGTGACGCTGACAGGCCATCACAATGCACGGATGAGCCTCGCTGAATGTCTGACGGTGATGAAGAAGGCCAAGCAGAGCCTTCACCAGTCTGGCCCATGGAAGCGCCATGTTGGCCCGGTGGTCGAAGGGCACGTCACGGCAACGGAGGTCAATCGGTCACGCCGCCATGGCTGGCACGTCCATTTTCACGTCCTGATGCTGATTGAGGCCGCCAGCGAGGCCGAAGCGATTGCGCTCGTGACCCGACACCTCGAACCTGAATGGCTCCGGCAGCTTGAGCGCCAAGGCAGCTGGGGCGGCGAACATGCGTTAGACGTCCAGGGCGCAGCCGATGCGGATCGGTATGTGGCCAAGTGGGGCGCGGCTGAGGAATTGGCTCTGTCGGGCGAAAAGCGGGCGCGCGGCGACGCCGAAATTAAGGGTCAAACGCCGTGGGAATTGTTGGCCGCTGCGGCTGATGGTGACGTCCAGGCTGGCCGCGATTTTGTGGAGTACGCGACCCTGTTTCATGGCAAACGCCAGCTCGTTTGGTCGCGAGGACTGAAAGAGCGTGTCGGCCTCAAGGTGGTGGAAGACGACGAGGCCGCCGGCGAGCTGGAGGATGTCGCAGATAGCGCCATCAACGAAGTGGCCGTGTTGCCGAGGCCGACTTGGCGTCACATCGTCCGGCGCAGGCTCCGCGCGGATCTGCTGGCCGCCGCTCGTGATGACGGATTCGATGGCGTCGCGCGGCTGCTCAATGAGCATGGCATCGGCGGAGCTATGCCCGGCGGCTGGCGACCAATGCTTAGCGAATATACCGAGCATATTGCGATGGCTGCGTGACTCTTTTGGCCCACCGGCCTGAGGGAAATTTGCCTCGCAGAGCTCACGATAATTTGACAAGGGCCGTAGGCCCGCAGGCAAATTGTCTAGTGAGTGTACAATTTTCAAATTCGAAGGTTATTGTAGAGCTGACGCTCTCAAGGTGATGACGCCTATGGCTGACGATTCTGGCGTCCGGCGGATTGTTTGTCGGATTAGCAAAATCCACACGATGGGCGTGCTGAAGGCCCGGTCGAATCACAATCTCCGAACGCCAGGTCATGACCAGCTCGCACATGTTGACCAGGCCAAATCCGCTCAGAATGAAATTCTGATTGGCACCGGCGACGCCATGCTCGATGTGCAATCAGCATTGAAATCGCGGCTGGTCGGCAACATTCGAAAAAATGGCGTTATCGCAGTGGAAATGGTGATGACAGCCCGGGCGGACTGGTTCGCTGCCGATCCCGCGCGAGCTGCCGCATTTAAGCAGCGGGTGCTGGACGAGGCGAATGCCAGATATGGGGCGAATCTCGTAACCGCAGTGTGGCACATGGATGAAGAGGCTCCCCATCTGCATCTGTATATCGTGCCGATGGACCAACGGGGCAAACTGAACTGCCGCGCGCAGTTCTCGACGATGGCGTCGCTGAAACAACTCCAGGATTGGGCTGGCGTGGTCGGCAATCCACTCGGCCTCGAACGCGGACGGCCAAAAGTGGAGTTTATTGTCGATGGCGATGAGCCGCCGGAGAACCAAAGCCCGGCGGAATATCGAGCTCAAACCAAACGCGACGCTGAAGCGGCAGCCGCCGCGCGGATCGCTGCCGAGCGTGATGCTGCCGAGGCTGCGCGACTGCTGATTGAGGCTCAGGAGCAGGCCGCCGCGATCCGCGCGGCCCAGGCTGAAATTATGGCAGAGCGTCGCCAGCTCAATGAGGATAAGGCGAGTTTTGCTGAGACGCTGCGTAGCCACACGAGGGCGATGGGCCTGGCGTTTCGTTCATGGCTGGCCAATTTCGTCGGCCGCGGCCAGATGCAGGCTGATGGAGGGCCATTGCTGACCTTTAAAGCCGCATGCCCCGCCAGTAAGCGAGAGGAGCTACAGTCCGAGCCAGCTAATGCGTACGCGCGCCTGGTGGTCGCTCATATGCCTGATCGTGACGCTCTTGCCGTGCTCGAAACCAAAAGCGCGGAGGCCAAGGAATAGCTTGGCGTAATCATGGGAAGATACTTAACAGGGAAGTGAGAGGGCCGCGGCAAAGCCGTTTTTCCATAGGCTCCGCCCCCCTGACAAGCATCACGAAATCTGACGCTCAAATCAGTGGTGGCGAAACCCGACAGGACTATAAAGATACCAGGCGTTTCCCCCTGGCGGCTCCCTCGTGCGCTCTCCTGTTCCTGCCTTTCGGTTTACCGGTGTCATTCCGCTGTTATGGCCGCGTTTGTCTCATTCCACGCCTGACACTCAGTTCCGGGTAGGCAGTTCGCTCCAAGCTGGACTGTATGCACGAACCCCCCGTTCAGTCCGACCGCTGCGCCTTATCCGGTAACTATCGTCTTGAGTCCAACCCGGAAAGACATGCAAAAGCACCACTGGCAGCAGCCACTGGTAATTGATTTAGAGGAGTTAGTCTTGAAGTCATGCGCCGGTTAAGGCTAAACTGAAAGGACAAGTTTTGGTGACTGCGCTCCTCCAAGCCAGTTACCTCGGTTCAAAGAGTTGGTAGCTCAGAGAACCTTCGAAAAACCGCCCTGCAAGGCGGTTTTTTCGTTTTCAGAGCAAGAGATTACGCGCAGACCAAAACGATCTCAAGAAGATCATCTTATTAATCAGATAAAATATTTCAAGATTTAGAACTCCAGCATGAGATCCCCGCGCTGGAGGATCATCCAGCCGGCGTCCCGGAAAACGATTCCGAAGCCCAACCTTTCATAGAAGGCGGCGGTGGAATCGAAATCTCGTGATGGCAGGTTGGGCGTCGCTTGGTCGGTCATTTCGAACCCCAGAGTCCCGCTCAGAAGAACTCGTCAAGAAGGCGATAGAAGGCGATGCGCTGCGAATCGGGAGCGGCGATACCGTAAAGCACGAGGAAGCGGTCAGCCCATTCGCCGCCAAGCTCTTCAGCAATATCACGGGTAGCCAACGCTATGTCCTGATAGCGGTCCGCCACACCCAGCCGGCCACAGTCGATGAATCCAGAAAAGCGGCCATTTTCCACCATGATATTCGGCAAGCAGGCATCGCCATGGGTCACGACGAGATCCTCGCCGTCGGGCATGCGCGCCTTGAGCCTGGCGAACAGTTCGGCTGGCGCGAGCCCCTGATGCTCTTCGTCCAGATCATCCTGATCGACAAGACCGGCTTCCATCCGAGTACGTGCTCGCTCGATGCGATGTTTCGCTTGGTGGTCGAATGGGCAGGTAGCCGGATCAAGCGTATGCAGCCGCCGCATTGCATCAGCCATGATGGATACTTTCTCGGCAGGAGCAAGGTGAGATGACAGGAGATCCTGCCCCGGCACTTCGCCCAATAGCAGCCAGTCCCTTCCCGCTTCAGTGACAACGTCGAGCACAGCTGCGCAAGGAACGCCCGTCGTGGCCAGCCACGATAGCCGCGCTGCCTCGTCCTGCAGTTCATTCAGGGCACCGGACAGGTCGGTCTTGACAAAAAGAACCGGGCGCCCCTGCGCTGACAGCCGGAACACGGCGGCATCAGAGCAGCCGATTGTCTGTTGTGCCCAGTCATAGCCGAATAGCCTCTCCACCCAAGCGGCCGGAGAACCTGCGTGCAATCCATCTTGTTCAATCATGCGAAACGATCCTCATCCTGTCTCTTGATCAGATCTTGATCCCCTGCGCCATCAGATCCTTGGCGGCAAGAAAGCCATCCAGTTTACTTTGCAGGGCTTCCCAACCTTACCAGAGGGCGCCCCAGCTGGCAATTCCGGTTCGCTTGCTGTCCATAAAACCGCCCAGTCTAGCTATCGCCATGTAAGCCCACTGCAAGCTACCTGCTTTCTCTTTGCGCTTGCGTTTTCCCTTGTCCAGATAGCCCAGGGGAAGCGTGGCGCTTTTCCGCTGCATAACCCTGCTTCGGGGTCATTATAGCGATTTTTTCGGTATATCCATCCTTTTTCGCACGATATACAGGATTTTGCCAAAGGGTTCGTGTAGACTTTCCTTGGTGTATCCAACGGCGTCAGCCGGGCAGGATAGGTGAAGTAGGCCCACCCGCGAGCGGGTGTTCCTTCTTCACTGTCCCTTATTCGCACCTGGCGGTGCTCAACGGGAATCCTGCTCTGCGAGGCTGGCCGGCTACCGCCGGCGTAACAGATGAGGGCAAGCGGATGGCTGATGAAACCAAGCCAACCAGGAAGGGCAGCCCACCTATCAAGGTGTACTGCCTTCCAGACGAACGAAGAGCGATTGAGGAAAAGGCGGCGGCGGCCGGCATGAGCCTGTCGGCCTACCTGCTGGCCGTCGGCCAGGGCTACAAAATCACGGGCGTCGTGGACTATGAGCACGTCCGCGAGCTGGCCCGCATCAATGGCGACCTGGGCCGCCTGGGCGGCCTGCTGAAACTCTGGCTCACCGACGACCCGCGCACGGCGCGGTTCGGTGATGCCACGATCCTCGCCCTGCTGGCGAAGATCGAAGAGAAGCAGGACGAGCTTGGCAAGGTCATGATGGGCGTGGTCCGCCCGAGGGCAGAGCCATGACTTTTTTAGCCGCTAAAACGGCCGGGGGGTGCGCGTGATTGCCAAGCACGTCCCCATGCGCTCCATACAGTTAGGGCGCGCCCAGCTG |
| Ptac/LacI-mamC^a^ | TCTAGGGCGGCGGATTTGTCCTACTCAGGAGAGCGTTCACCGACAAACAACAGATAAAACGAAAGGCCCAGTCTTTCGACTGAGCCTTTCGTTTTATTTGATGCCTTTAATTAAAGCGGATAACAATTTCAGAATTCGCGCTCACTGCCCGCTTTCCAGTCGGGAAACCTGTCGTGCCAGCTGCATTAATGAATCGGCCAACGCGCGGGGAGAGGCGGTTTGCGTATTGGGCGCCAGGGTGGTTTTTCTTTTCACCAGTGAGACGGGCAACAGCTGATTGCCCTTCACCGCCTGGCCCTGAGAGAGTTGCAGCAAGCGGTCCACGCTGGTTTGCCCCAGCAGGCGAAAATCCTGTTTGATGGTGGTTAACGGCGGGATATAACATGAGCTGTCTTCGGTATCGTCGTATCCCACTACCGAGATATCCGCACCAACGCGCAGCCCGGACTCGGTAATGGCGCGCATTGCGCCCAGCGCCATCTGATCGTTGGCAACCAGCATCGCAGTGGGAACGATGCCCTCATTCAGCATTTGCATGGTTTGTTGAAAACCGGACATGGCACTCCAGTCGCCTTCCCGTTCCGCTATCGGCTGAATTTGATTGCGAGTGAGATATTTATGCCAGCCAGCCAGACGCAGACGCGCCGAGACAGAACTTAATGGGCCCGCTAACAGCGCGATTTGCTGGTGACCCAATGCGACCAGATGCTCCACGCCCAGTCGCGTACCGTCTTCATGGGAGAAAATAATACTGTTGATGGGTGTCTGGTCAGAGACATCAAGAAATAACGCCGGAACATTAGTGCAGGCAGCTTCCACAGCAATGGCATCCTGGTCATCCAGCGGATAGTTAATGATCAGCCCACTGACGCGTTGCGCGAGAAGATTGTGCACCGCCGCTTTACAGGCTTCGACGCCGCTTCGTTCTACCATCGACACCACCACGCTGGCACCCAGTTGATCGGCGCGAGATTTAATCGCCGCGACAATTTGCGACGGCGCGTGCAGGGCCAGACTGGAGGTGGCAACGCCAATCAGCAACGACTGTTTGCCCGCCAGTTGTTGTGCCACGCGGTTGGGAATGTAATTCAGCTCCGCCATCGCCGCTTCCACTTTTTCCCGCGTTTTCGCAGAAACGTGGCTGGCCTGGTTCACCACGCGGGAAACGGTCTGATAAGAGACACCGGCATACTCTGCGACATCGTATAACGTTACTGGTTTCACATTCACCACCCTGAATTGACTCTCTTCCGGGCGCTATCATGCCATACCGCGAAAGGTTTTGCACCATTCGATGGTGTCAACGTAAATGCATGCCGCTTCGCCTTCGCGCGCGAATTGCAGGTACCATTTATCAGGGTTATTGTCTCATGAGCGGATACATATTTGAATGTATTTAGAAAAATAAACAAATAGGGGTTCCGCGCACATTTCCCCGAAAAGTGCCACCTGACGTCTAAGAAACCATTATTATCATGACATTAACCTATAAAAATAGGCGTATCACGAGGCCCTTTCGTCTTCACCTCGAGAAAATTTATCAAAAAGAGTGTTGACTTGTGAGCGGATAACAATGATACTTAGATTCAATTGTGAGCGGATAACAATTTCACACAATCGATAGCTGTCACCGGATGTGCTTTCCGGTCTGATGAGTCCGTGAGGACGAAACAGCCTCTACAAATAATTTTGTTTAATCTAGAAATAATTTTGTTTATCTCTCGAGGAGGATTCGCCATGCCCTTTCACCTTGCCCCCTATCTGGCGAAATCCGTTCCCGGCGTCGGCGTTCTCGGCGCCCTGGTCGGCGGCGCCGCCGCCTTGGCCAAGAACGTCCGCCTCCTGAAGGAAAAGCGCATCACCAATACCGAAGCGGCCATCGATACCGGCAAGGAAACCGTCGGCGCCGGCCTGGCCACCGCGCTTTCCGCCGTGGCCGCGACCGCCGTCGGCGGCGGCCTGGTTGTATCGCTGGGCACCGCCTTGGTGGCCGGCGTTGCCGCCAAATATGCCTGGGATCGCGGCGTCGATCTGGTCGAGAAGGAACTGAACCGCGGCAAAGCTGCCAACGGCGCTTCCGACGAGGACATCTTGCGGGACGAACTGGCCTGATAAGCTTGCGGCCGCGTCGTGACTGGGAAAACCCTGGCGACTAGTCTTGGACTCCTGTTGATAGATCCAGTAATGACCTCAGAACTCCATCTGGATTTGTTCAGAACGCTCGGTTGCCGCCGGGCGTTTTTTATTGGTGAGAATCCAG |

^a^DNA sequence colors corresponds to terminators (red), spacers (black), lacI-CDS (blue), promoter (green), ribozyme insulators (purple), and mamC-CDS (orange).

**Supplementary Table 2. DNA sequence of the mamC-peptide CDS**

| **CDS** | **DNA Sequence^a^** |
| --- | --- |
| mamC-his | ATGCCCTTTCACCTTGCCCCCTATCTGGCGAAATCCGTTCCCGGCGTCGGCGTTCTCGGCGCCCTGGTCGGCGGCGCCGCCGCCTTGGCCAAGAACGTCCGCCTCCTGAAGGAAAAGCGCATCACCAATACCGAAGCGGCCATCGATACCGGCAAGGAAACCGTCGGCGCCGGCCTGGCCACCGCGCTTTCCGCCGTGGCCGCGACCGCCGTCGGCGGCGGCCTGGTTGTATCGCTGGGCACCGCCTTGGTGGCCGGCGTTGCCGCCAAATATGCCTGGGATCGCGGCGTCGATCTGGTCGAGAAGGAACTGAACCGCGGCAAAGCTGCCAACGGCGCTTCCGACGAGGACATCTTGCGGGACGAACTGGCCGCTGGTGGATCCCATCACCATCACCATCACCATCACCATTGA |
| mamC-pHLIP | ATGCCCTTTCACCTTGCCCCCTATCTGGCGAAATCCGTTCCCGGCGTCGGCGTTCTCGGCGCCCTGGTCGGCGGCGCCGCCGCCTTGGCCAAGAACGTCCGCCTCCTGAAGGAAAAGCGCATCACCAATACCGAAGCGGCCATCGATACCGGCAAGGAAACCGTCGGCGCCGGCCTGGCCACCGCGCTTTCCGCCGTGGCCGCGACCGCCGTCGGCGGCGGCCTGGTTGTATCGCTGGGCACCGCCTTGGTGGCCGGCGTTGCCGCCAAATATGCCTGGGATCGCGGCGTCGATCTGGTCGAGAAGGAACTGAACCGCGGCAAAGCTGCCAACGGCGCTTCCGACGAGGACATCTTGCGGGACGAACTGGCCGCTGCGGAACAGAACCCGATTTATTGGGCGCGCTATGCGGATTGGCTGTTTACCACCCCGCTGCTCCTGCTGGATCTGGCGCTCCTGGTGGATGCGGATGAAGGCACCTGA |
| mamC-iRGD | ATGCCCTTTCACCTTGCCCCCTATCTGGCGAAATCCGTTCCCGGCGTCGGCGTTCTCGGCGCCCTGGTCGGCGGCGCCGCCGCCTTGGCCAAGAACGTCCGCCTCCTGAAGGAAAAGCGCATCACCAATACCGAAGCGGCCATCGATACCGGCAAGGAAACCGTCGGCGCCGGCCTGGCCACCGCGCTTTCCGCCGTGGCCGCGACCGCCGTCGGCGGCGGCCTGGTTGTATCGCTGGGCACCGCCTTGGTGGCCGGCGTTGCCGCCAAATATGCCTGGGATCGCGGCGTCGATCTGGTCGAGAAGGAACTGAACCGCGGCAAAGCTGCCAACGGCGCTTCCGACGAGGACATCTTGCGGGACGAACTGGCCGCTGGTGGATCCTGCCGTGGCGACAAGGGTCCGGATTGCTGA |

^a^DNA sequence colors corresponds to mamC (orange), AGGS linker (blue), and peptide tag (green).
